# Heterotrimeric kinesin-2 autoinhibition mediated by interactions of the CC2 and proximal tail domains with the motor domains is essential for cilium formation and maintenance

**DOI:** 10.64898/2026.05.19.726381

**Authors:** Jessica M. Adams, Caleb Sawe, Martin F. Engelke

**Affiliations:** Department of Biochemistry & Cellular and Molecular Biology, University of Tennessee, Knoxville, TN, USA; School of Biological Sciences, Illinois State University, Normal, IL, USA

## Abstract

Autoinhibition is a fundamental regulatory mechanism for kinesins, including the heterotrimeric kinesin-2 complex (KIF3A/KIF3B/KAP3), which mediates cytoplasmic cargo transport and anterograde intraflagellar transport. In mammals, kinesin-2 is essential for ciliogenesis and ciliary function. Although a structural model of autoinhibited kinesin-2 has been proposed, further validation and functional analysis are needed. Here, we elucidate the mechanism and functional importance of kinesin-2 autoinhibition using cell-based assays guided by structural predictions. Through knockout-rescue experiments with chimeric KIF3A-KIF3B subunits, we show that subunit-specific interactions between the motor domains and the C-terminal coiled-coil domains and adjacent tail β-hairpin motifs stabilize the autoinhibited state. Interestingly, these same C-terminal regions required for autoinhibition are required for stable heterodimerization of the motor. We further find that a flexible region within the coiled-coil stalk is required for this autoregulation and may facilitate the underlying conformation transitions. Furthermore, the capacity to autoinhibit directly correlates with the ability of mutant motors to support ciliogenesis, underscoring the significance of autoinhibition for motor function. Collectively, these findings define key subunit-specific interactions underlying kinesin-2 autoinhibition, identify elements that govern conformational transitions, and demonstrate that autoinhibition is essential for kinesin-2 function in ciliogenesis.

## INTRODUCTION

Precise spatiotemporal control of intracellular transport is essential for cellular organization. In transport kinesins, and likely more broadly across the kinesin superfamily, autoinhibition is a central mechanism of this control (Verhey and Hammond, 2009). Kinesins are thought to reside in a compact, backfolded inactive state until cellular cues, including cargo binding, promote transition to an extended, active conformation. In kinesin-1, this inactive state is mediated by folding around a central elbow that functions as a hinge, enabling inhibitory tail-motor interactions that suppress ATPase activity and microtubule binding (Coy et al., 1999; Friedman and Vale, 1999; Hackney and Stock, 2000; Tan et al., 2023; Weijman et al., 2022). Although the mechanism of kinesin-1 autoinhibition has been studied extensively, how autoinhibition is achieved in other kinesin families remains less well understood (Tan et al., 2026; Yildiz, 2025). This question is important because loss of autoinhibition can have substantial physiological consequences: experimentally relieving autoinhibition produces unphysiological motor activity and altered cargo distribution in cells (Kelliher et al., 2018), and disease-associated mutations that disrupt autoinhibition of KIF5A and KIF21A have been linked to neurological dysfunction (Baron et al., 2022; Cheng et al., 2014).

In mammals, the kinesin-2 family comprises four motor subunits: KIF3A, KIF3B, KIF3C, and KIF17. While KIF17 functions as a homodimer, the canonical kinesin-2 motors are formed by KIF3A heterodimerization with either KIF3B or KIF3C. These heterodimers associate with the nonmotor accessory subunit KAP3, making kinesin-2 unique within the kinesin superfamily in containing the only well-established heterotrimeric kinesins (Scholey, 2013). Among these, the KIF3A/KIF3B/KAP3 complex is the sole anterograde intraflagellar transport (IFT) motor in mammals and is therefore essential for cilium assembly, maintenance, and signaling (Engelke et al., 2019). While KAP3 primarily mediates cargo binding, the spatiotemporal regulation of this complex largely depends on the autoinhibition of its motor subunits, which is the focus of this study.

Structurally, KIF3A and KIF3B each contain an N-terminal motor domain, a central α-helical coiled-coil stalk that mediates heterodimerization, and a divergent C-terminal tail domain (**Fig. 1A**). Early electron microscopy studies showed that sea urchin kinesin-2 can adopt both extended and folded conformations, suggesting that the conformational switching characteristic of kinesin-1 could also apply to kinesin-2 (Wedaman et al., 1996). More recently, an AlphaFold3-generated model that fit a single-particle negative-stain electron microscopy density of the autoinhibited kinesin-2 complex provided an initial structural view of this folded architecture (Webb et al., 2025). This model suggested that the stalk collapses onto the motor domains rather than undergoing a simple hinge-like backfolding reaction, as described for kinesin-1, and identified a novel β-hairpin motif within the tail domain that mediates critical motor-tail contacts, stabilizing the autoinhibited state. However, the broader structural requirements that promote and stabilize this asymmetric folded conformation have not been extensively tested experimentally in cells, and the extent to which this regulatory mechanism governs ciliary function remains incompletely understood.

**Figure 1.**
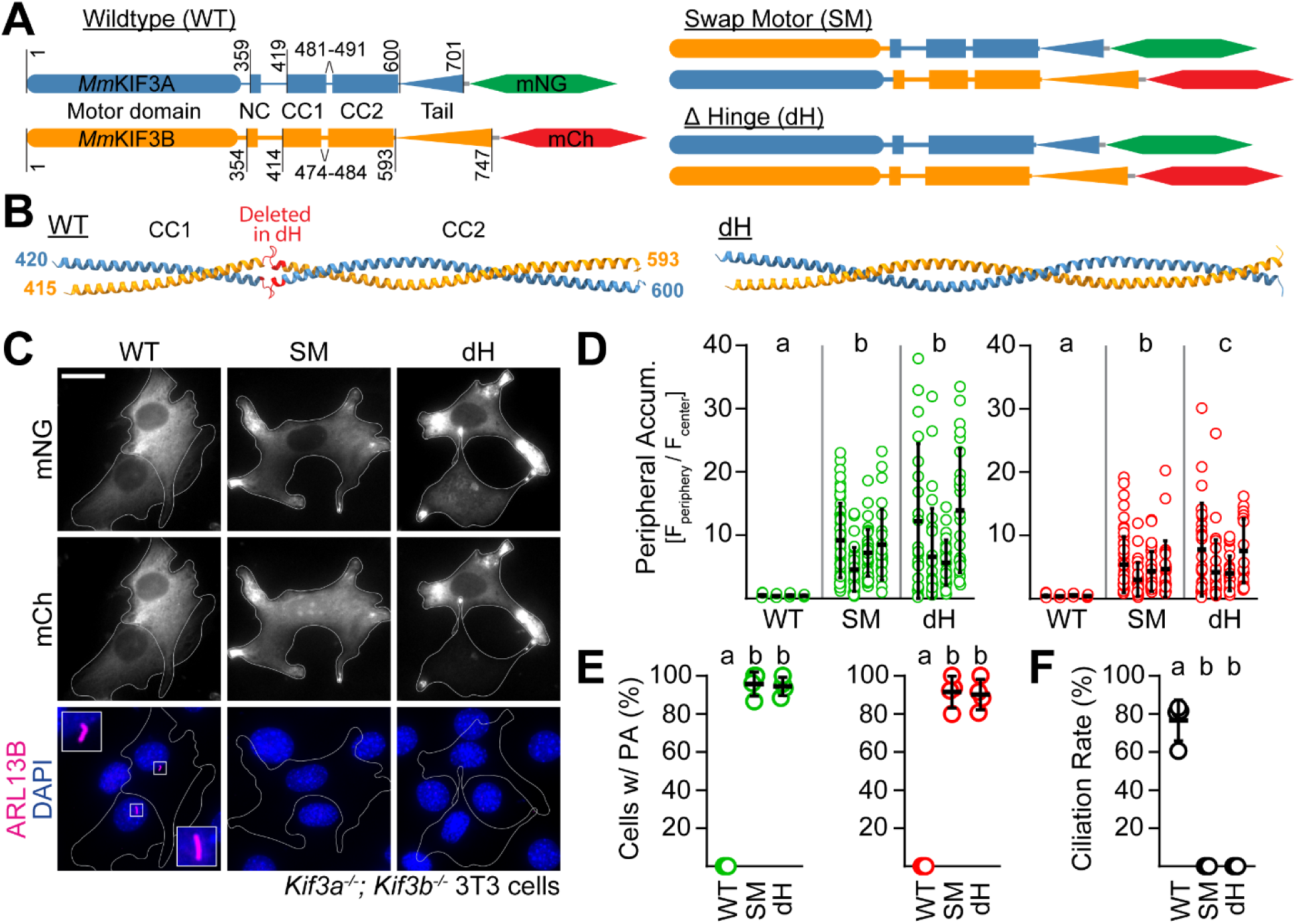
A flexible hinge region in the KIF3A/KIF3B stalk is required for autoinhibition and ciliogenesis. **(A)** To-scale schematic of the *Mm*KIF3A (blue) and *Mm*KIF3B (orange) proteins with the motor domains, neck coil (NC), coiled-coil 1 (CC1), coiled-coil 2 (CC2), and tail domains annotated. Proteins were fused to the C-terminus of the fluorescent proteins mNeonGreen (mNG) or mCherry (mCh) via a short linker. Amino acid residues at which splicing occurred to generate the Swap Motor (SM) and Δ Hinge (dH) constructs (right), as well as residues used to model the stalk region (panel B), are indicated. Full sequences of translated proteins are available in Supplementary Table 1. **(B)** AlphaFold 3-predicted structure of the CC1 and CC2 domains of WT KIF3A/KIF3B stalk (left). Residues comprising the flexible hinge were identified and removed to generate a predicted continuous coiled-coil while preserving heptad register (right). **(C-F)** *Kif3a^-/-^; Kif3b^-/-^* 3T3 cells were co-transfected with two expression plasmids encoding the pairs of proteins illustrated in panel A, and primary cilium formation was stimulated by serum starvation for 48 hours. **(C)** Representative image of each condition. Cilia were visualized by immunostaining with an antibody against the ciliary membrane marker ARL13B (magenta), and nuclei were visualized with DAPI (blue). Insets show an enlarged region of interest containing the primary cilium. Scale bar = 20 μm. **(D)** Quantification of peripheral accumulation (PA) ratio (fluorescence intensity of a peripheral [F_periphery_] divided by a perinuclear [F_center_] region of interest) in transfected cells in each condition (mNG, green, left; mCh, red, right). **(E)** Quantification of the percentage of cells in each condition with PA ratios greater than 1 (mNG, green, left; mCh, red, right). **(F)** Quantification of the ciliation rate (percentage of transfected cells with a cilium) in each condition. Data represent four biological replicates, with 24-68 cells analyzed per condition per replicate. Data are presented as mean ± SD and were analyzed via one-way ANOVA followed by Tukey’s post hoc tests. Groups that do not share a letter differ significantly (compact letter display). All summary statistics, ANOVA outputs, and Tukey post hoc results are provided in Supplementary Tables 2-4.

Here, we build on this structural framework to refine the mechanism of mammalian heterotrimeric kinesin-2 autoinhibition and test its importance for ciliogenesis. We find that, as proposed for other kinesin-2 motors (Brunnbauer et al., 2010; Hammond et al., 2010; Imanishi et al., 2006), a critical diglycine hinge within the coiled-coil stalk is required for autoinhibition and likely enables the motor to transition into the compact, collapsed conformation. We initially sought to define the domains required for autoinhibition using conventional truncation strategies. These experiments revealed an important complication: in addition to the previously recognized role of the second coiled-coil segment (CC2) in heterodimerization (De Marco et al., 2001; Vukajlovic et al., 2011), the proximal tail regions also contributed to the stable assembly of KIF3A/KIF3B. We therefore adopted a targeted domain-swapping strategy to map the core autoinhibitory elements. Our cell-based analyses confirm the critical role of the tail β-hairpin and the second coiled-coil segment (CC2) in autoinhibition, while placing greater emphasis on the significance of CC2. We further uncover functional asymmetry between the two motor subunits, with the KIF3A motor domain contributing more strongly to autoinhibition than the KIF3B motor domain. Finally, we show that precise tuning of autoinhibition is essential for kinesin-2 function. Partial relief of autoinhibition yields hyperactive motors that still support ciliogenesis, albeit to a reduced extent, and accumulate at the ciliary tip. Together, our findings define a mechanistic framework for kinesin-2 autoinhibition and establish that this regulation is critical for ciliary assembly and maintenance.

## RESULTS

### A hinge in the coiled-coil stalk is required for autoinhibition

Transport kinesins are commonly autoinhibited through intramolecular interactions between the C-terminal cargo-binding region and the motor domain (Verhey and Hammond, 2009; Yildiz, 2025). Coiled-coil prediction (CoCoNat, Madeo et al., 2023) and AlphaFold 3 modeling (Abramson et al., 2024) of the KIF3A/KIF3B stalk identified two coiled-coil segments (CC1 and CC2) interrupted by a conserved di-glycine motif that could function as a hinge to enable motor-tail contact, as previously reported (**Fig. 1A**, Brunnbauer et al., 2010; Imanishi et al., 2006; Wedaman et al., 1996). Using aforementioned tools, we generated a construct in which residues encompassing the flexible hinge were deleted, and CC1 and CC2 were fused while preserving the native heptad register (dH, **Fig. 1B**). When expressed in *Kif3a^-/-^; Kif3b^-/-^* mouse embryonic fibroblast 3T3 cells, C-terminally fluorescently-tagged wildtype (WT) kinesin-2 displayed the homogeneous, diffuse cytoplasmic distribution characteristic of autoinhibited kinesins (**Fig. 1C**, Cai et al., 2007; Hammond et al., 2010; Hollenbeck, 1989). By contrast and despite low expression, the dH mutant motor strongly accumulated in the cell periphery, similarly to a previously reported chimeric motor that was constitutively activated by fusing the KIF3A motor domain to the KIF3B stalk and tail domains and vice versa (Swap Motor; SM), used here as a positive control (Adams et al., 2024; Brunnbauer et al., 2010). Quantification of peripheral accumulation as the ratio of peripheral to perinuclear fluorescence intensity showed that SM and dH motors were strongly activated to a similar extent (**Fig. 1D**). Consistent with this, most cells transfected with SM and dH constructs exhibited peripheral accumulation (defined as a ratio > 1), whereas cells expressing the WT constructs rarely exceeded this threshold (**Fig. 1E**). *Kif3a^-/-^; Kif3b^-/-^* cells cannot make cilia, as they lack the anterograde intraflagellar transport (IFT) motor (Engelke et al., 2019). Re-expression of WT KIF3A and KIF3B rescued ciliation in these cells to WT levels, as expected (**Fig. 1F**, Adams et al., 2024). In contrast, re-expression of SM and dH constructs did not rescue ciliogenesis. Together, these results indicate that the hinge in the central stalk is required for kinesin-2 to adopt the autoinhibited state and that loss of autoinhibition compromises heterotrimeric kinesin-2 function during ciliogenesis, consistent with prior work (Webb et al., 2025).

### Truncation of kinesin-2 disrupts dimerization

We next sought to identify which regions of the C-terminal cargo-binding portion contribute to autoinhibition. To this end, we generated a series of KIF3A and KIF3B truncations, a strategy that has been successfully used to study autoinhibition in other kinesins (Coy et al., 1999; Friedman and Vale, 1999; Hammond et al., 2010). Since bulky tags, especially on the tail domain, have been reported to affect motor activity (Fasawe et al., 2024; Hammond et al., 2010), we N-terminally tagged the KIF subunits with myc-mNG or HA-mCh for this assay (**Fig. 2A**). The change of tagging location did not significantly affect the peripheral accumulation of the wildtype (WT-N) motor (**Fig. 2B-D**), nor did it impact the ciliation rate of transfected cells following serum starvation, which was similar to previous assays at 79.7±4.4%. Surprisingly, repositioning the tag from the C- to the N-terminus significantly reduced the peripheral accumulation ratio and the number of cells exhibiting peripheral accumulation in cells expressing the Swap Motor (SM) constructs. This reduction in motor activation might be due to the N-terminal tag interfering with motor domain function or to C-terminal tagging mimicking cargo, thereby activating the motor. The former is less likely, as WT motor function in ciliogenesis was not impaired by N-terminal tagging. Irrespective of the underlying cause, this tagging effect must be taken into account when interpreting results from peripheral accumulation assays. We also observed that peripheral accumulation measurements differed somewhat between the mNG and mCh channels. This likely results from differences in expression levels between the two constructs, as well as a tendency of the N-terminally HA-mCh Swap construct to localize to the nucleus, artificially inflating measurements in the perinuclear region.

**Figure 2.**
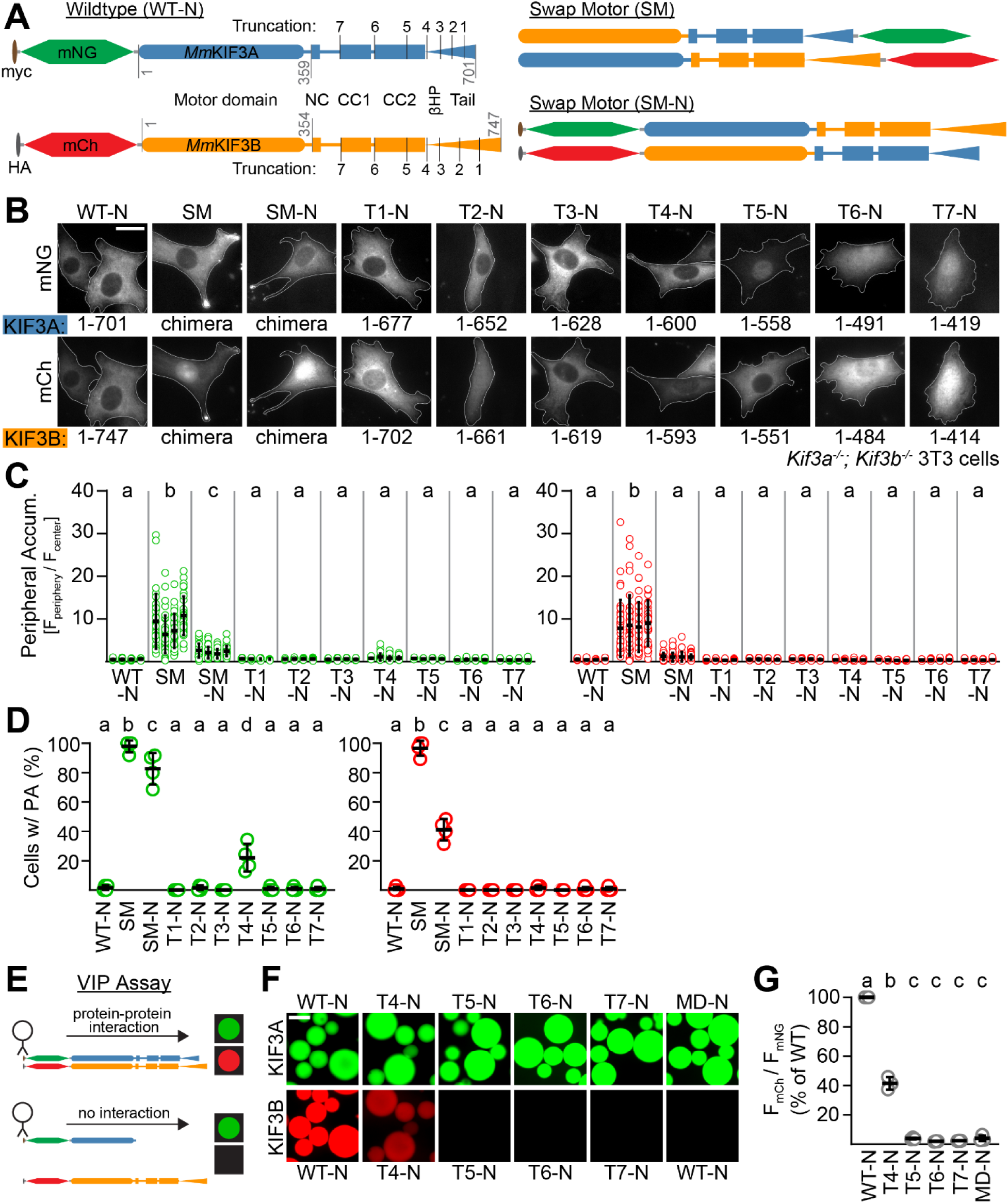
Truncation of KIF3A/KIF3B prevents motor dimerization. **(A)** Schematic of *Mm*KIF3A (blue) and *Mm*KIF3B (orange) proteins annotated as in Figure 1, with the β-hairpin (βHP) motif in the tail domains also labeled. Proteins (except for SM) were dual-tagged on the N-terminus with myc-mNG or HA-mCh via a short linker. Amino acid positions at which truncations were performed are indicated in black, and the residue numbers are given in Panel B. Residues at which splicing occurred to generate the Swap Motor constructs are indicated in gray. Full sequences of translated proteins are available in Supplementary Table 1. **(B-C)** *Kif3a^-/-^; Kif3b^-/-^*3T3 cells were co-transfected with two expression plasmids encoding the pairs of proteins illustrated in panel A and cultured for 48 hours. **(B)** Representative image of each condition. The residues of *Mm*KIF3A and *Mm*KIF3B included in each truncation are listed below each image, while the shorthand designation (e.g., T1 for Truncation 1) is labeled above. Scale bar = 20 μm. **(C)** Quantification of peripheral accumulation (PA) ratio (fluorescence intensity of a peripheral [F_periphery_] divided by a perinuclear [F_center_] region of interest) in transfected cells in each condition (mNG, green, left; mCh, red, right). **(D)** Quantification of the percentage of cells in each condition with PA ratios greater than 1 (mNG, green, left; mCh, red, right). **(E-G)** The Visible Immunoprecipitation (VIP) assay was used to evaluate the dimerization status of pairs of truncated motors. **(E)** Schematic of the VIP assay. Whole cell lysates from COS7 cells co-transfected with pairs of truncated motors were prepared and incubated with myc-antibody-coated agarose beads, which were then imaged via fluorescence microscopy. **(F)** Representative images of beads. Scale bar = 100 μm. **(G)** Quantification of the ratio of fluorescence intensity of mCh-KIF3B to that of mNG-KIF3A in each set of truncation mutants, normalized to that of WT in each replicate. Data represent four biological replicates (C-D) or three (G), with 30-37 cells (C,D) or 31-60 beads (G) analyzed per condition per replicate. Data are presented as mean ± SD and were analyzed via one-way ANOVA followed by Tukey’s post hoc tests. Groups that do not share a letter differ significantly (compact letter display). All summary statistics, ANOVA outputs, and Tukey post hoc results are provided in Supplementary Tables 2-4.

Next, we evaluated the peripheral accumulation phenotypes of the successively truncated motors, with T1 being the least and T7 the most truncated construct (**Fig. 2A**). Truncation of most of the tail domains (T1-T3) did not lead to motor activation, as judged by the degree to which motor accumulated as well as the number of cells showing peripheral accumulation (**Fig. 2B-D**). Interestingly, the removal of the entire tail domains (T4), including the recently discovered β-hairpin motifs (Jiang et al., 2025; Webb et al., 2025), led to modest but significant activation, as judged by the percentage of cells showing mNG accumulation (**Fig. 2D**). This activation was, however, not observed for mCh. The discrepancy between the two signals could be explained by a recent report that KIF3B subunits might form homodimers (Jiang et al., 2026), by decreased heterodimerization and appearance of monomeric species caused by tail loss, or by a combination of both. Further truncations that removed increasing segments of the coiled-coil stalk (T5-T7) showed no activation phenotype (**Fig. 2B-D**), consistent with the notion that the tail and coiled-coil domains promote dimerization.

To test this hypothesis, we used a simple protein interaction assay, the Visible Immunoprecipitation (VIP) assay (Katoh et al., 2018). Cell lysates from COS-7 cells co-expressing pairs of truncated constructs were incubated with anti-myc agarose beads, and associated fluorescence was visualized by epifluorescence microscopy (**Fig. 2E-F**). All myc-mNG-tagged KIF3A constructs were readily detected on the beads, and full-length HA-mCh-tagged KIF3B interacted strongly with WT KIF3A, whereas the KIF3A motor domain alone (MD, residues 1-359, negative control) did not interact with full-length KIF3B (**Fig. 2F-G**). Complete truncation of the tail domains (T4) reduced the interaction between KIF3A and KIF3B, and further truncation beyond the tails abolished detectable interaction. These data indicate that stable KIF3A/KIF3B heterodimerization requires not only the C-terminal stalk region, as previously reported (De Marco et al., 2001; Vukajlovic et al., 2011), but also proximal regions of the tail domains. They also show that progressive truncation is not a suitable strategy for mapping autoinhibitory elements in kinesin-2, because the same deletions that remove candidate regulatory regions also disrupt dimer assembly.

### Domain-specific interactions stabilize the autoinhibited state

Because the truncation approach was confounded by loss of heterodimerization, we next turned to a strategy that preserves overall motor architecture while selectively altering subunit identity. Autoinhibition in the KIF3A/KIF3B heterodimer is expected to depend on subunit-specific intramolecular contacts, because the two polypeptides diverge especially outside the motor domain. Pairwise alignment of mouse KIF3A (NP_032469.2) and KIF3B (NP_032470.3) using EMBOSS (Needleman-Wunsch algorithm; Madeira et al., 2024; Rice et al., 2000) showed 67.5% identity in the motor domains and only 37.1% in the coiled-coil stalk and 13.3% in the tail. To coarsely map which domains contribute to autoinhibition, we designed a series of C-terminally fluorescently tagged chimeric KIF3A/KIF3B constructs. These Swap constructs exchanged different regions between subunits: the entire coiled-coil stalk in Swap Coiled-coil (SC), the neck coil plus adjacent charged regions in Swap Neck coil (SN), and only the tail domains in Swap Tail (ST) (**Fig. 3A**). Expression of the previously used controls, wildtype (WT) and Swap Motor (SM) KIF3A/KIF3B, resulted in the expected diffuse cytoplasmic and strongly peripherally accumulated phenotypes, respectively (**Fig. 3B**). While the intracellular distribution of SC motors resembled that of SM, SN motors resembled the WT control. Quantification of the peripheral accumulation level confirmed that SM and SC motors were activated to similar extents, whereas SN motors were indistinguishable from WT (**Fig. 3C**). Consistent with this, nearly all cells transfected with SM and SC constructs showed peripheral accumulation, whereas cells transfected WT and SN constructs rarely did (**Fig. 3D**). ST motors showed an intermediate phenotype with only modestly elevated peripheral accumulation ratios that were not significantly different from WT, yet a significant fraction of ST-expressing cells crossed the peripheral accumulation threshold (**Fig. 3C and D**). The above data indicate that a portion of the coiled-coil stalk comprising CC1 and/or CC2, but not the neck coil, is critically required for kinesin-2 autoinhibition, likely through specific interactions with the motor domains, while the orientation of the tail domains is less important.

**Figure 3.**
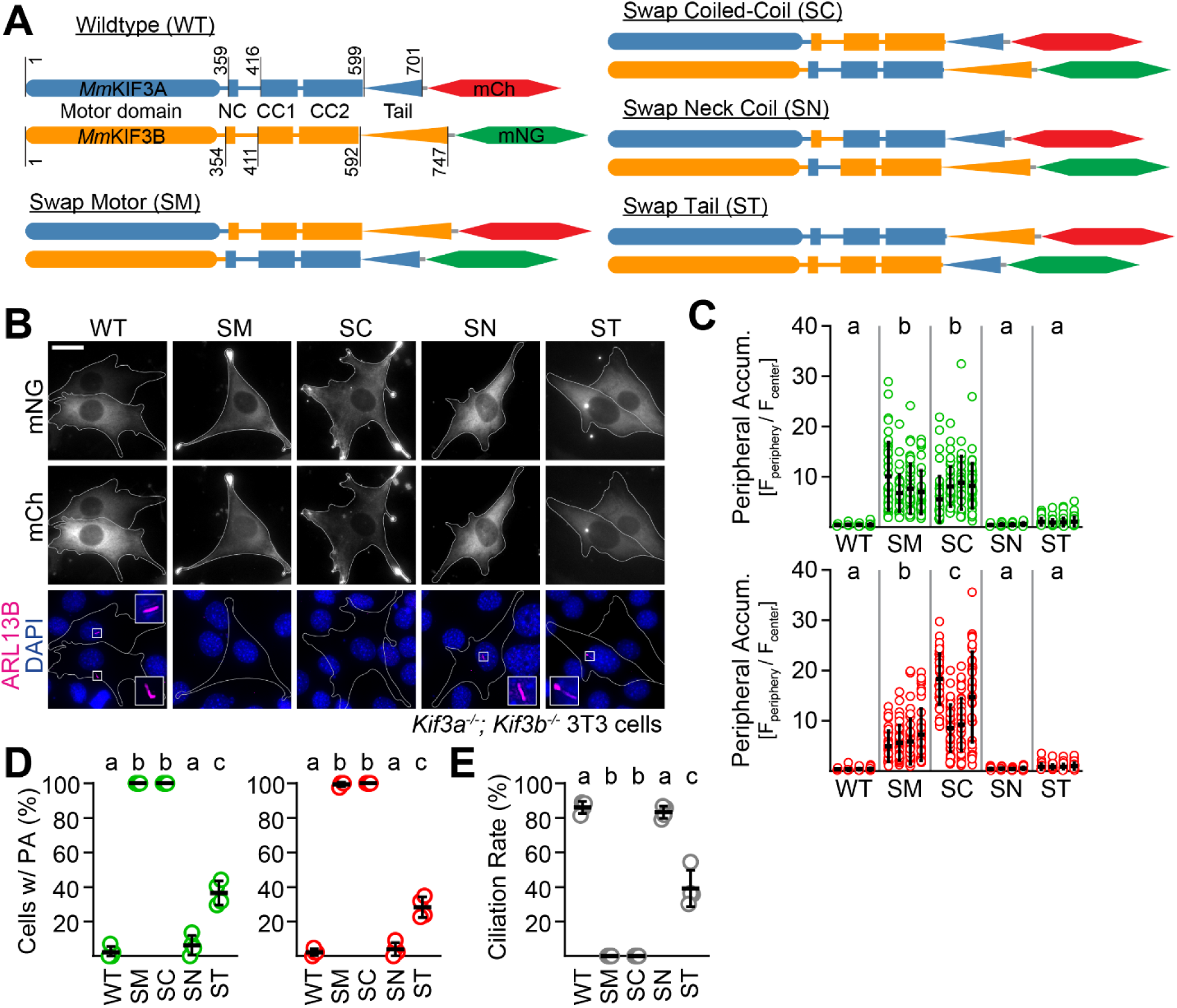
Domain-specific alterations disrupt autoinhibition of the heterodimeric KIF3A/KIF3B kinesin-2 motor. **(A)** Schematic of *Mm*KIF3A (blue) and *Mm*KIF3B (orange) proteins annotated as in Figure 1. Amino acid residues at which splicing occurred to generate the chimeric mutant motors are indicated. Full sequences of translated proteins are available in Supplementary Table 1. **(B-E)** *Kif3a^-/-^; Kif3b^-/-^* 3T3 cells were co-transfected with two expression plasmids encoding the pairs of proteins illustrated in panel A, and primary cilium formation was stimulated by serum starvation for 48 hours. **(B)** Representative image of each condition. Cilia were visualized by immunostaining with an antibody against the ciliary membrane marker ARL13B (magenta), and nuclei were visualized with DAPI. Insets show an enlarged region of interest containing the primary cilium. Scale bar = 20 μm. **(C)** Quantification of peripheral accumulation (PA) ratio (fluorescence intensity of a peripheral [F_periphery_] divided by a perinuclear [F_center_] region of interest) in transfected cells in each condition (mNG, green, top; mCh, red, bottom). **(D)** Quantification of the percentage of cells in each condition with PA ratios greater than 1 (mNG, green, left; mCh, red, right) **(E)** Quantification of the ciliation rate (percentage of transfected cells with a cilium) in each condition. Data represent four biological replicates, with 42-45 cells analyzed per condition per replicate. Data are presented as mean ± SD and were analyzed via one-way ANOVA followed by Tukey’s post hoc tests. Groups that do not share a letter differ significantly (compact letter display). All summary statistics, ANOVA outputs, and Tukey post hoc results are provided in Supplementary Tables 2-4.

Next, we examined the ability of the above constructs to rescue kinesin-2 function. We found that the SN constructs, which showed no peripheral accumulation, restored ciliation to high levels comparable to WT (**Fig. 3B and E**). In contrast, the SC constructs, which strongly accumulated at the periphery, did not rescue ciliogenesis, similar to the SM controls. The mild activation of ST supported ciliogenesis at a significantly reduced rate. Thus, the extent of autoinhibition loss correlates with the loss of kinesin-2 function, reinforcing the conclusion that autoinhibition is required for ciliogenesis.

### A previously identified β-hairpin tail domain structure and the adjacent CC2 domain both contribute substantially to autoinhibition

The domain-swap analysis implicated both the stalk and the tail domains in autoinhibition, but did not resolve which stalk segment is most important or whether specific tail elements are involved. In addition, our previous truncation assay was limited by the reduced sensitivity of N-terminally tagged constructs. We therefore revisited tail contributions using C-terminally tagged truncation mutants and, in parallel, subdivided the stalk into its two principal coiled-coil segments. Specifically, we generated C-terminally tagged T3 and T4 truncation constructs. T3 lacked most of the tail domain but retained the recently identified β-hairpin structures (Jiang et al., 2025; Webb et al., 2025), whereas T4 lacked the entire tail domain (**Fig. 4A**). To distinguish the contributions of the two coiled-coil stalk segments, we also generated Swap constructs in which either CC1 or CC2 was selectively swapped between KIF3A and KIF3B, enabling us to test the relative roles of CC1 and CC2 in autoinhibition.

**Figure 4.**
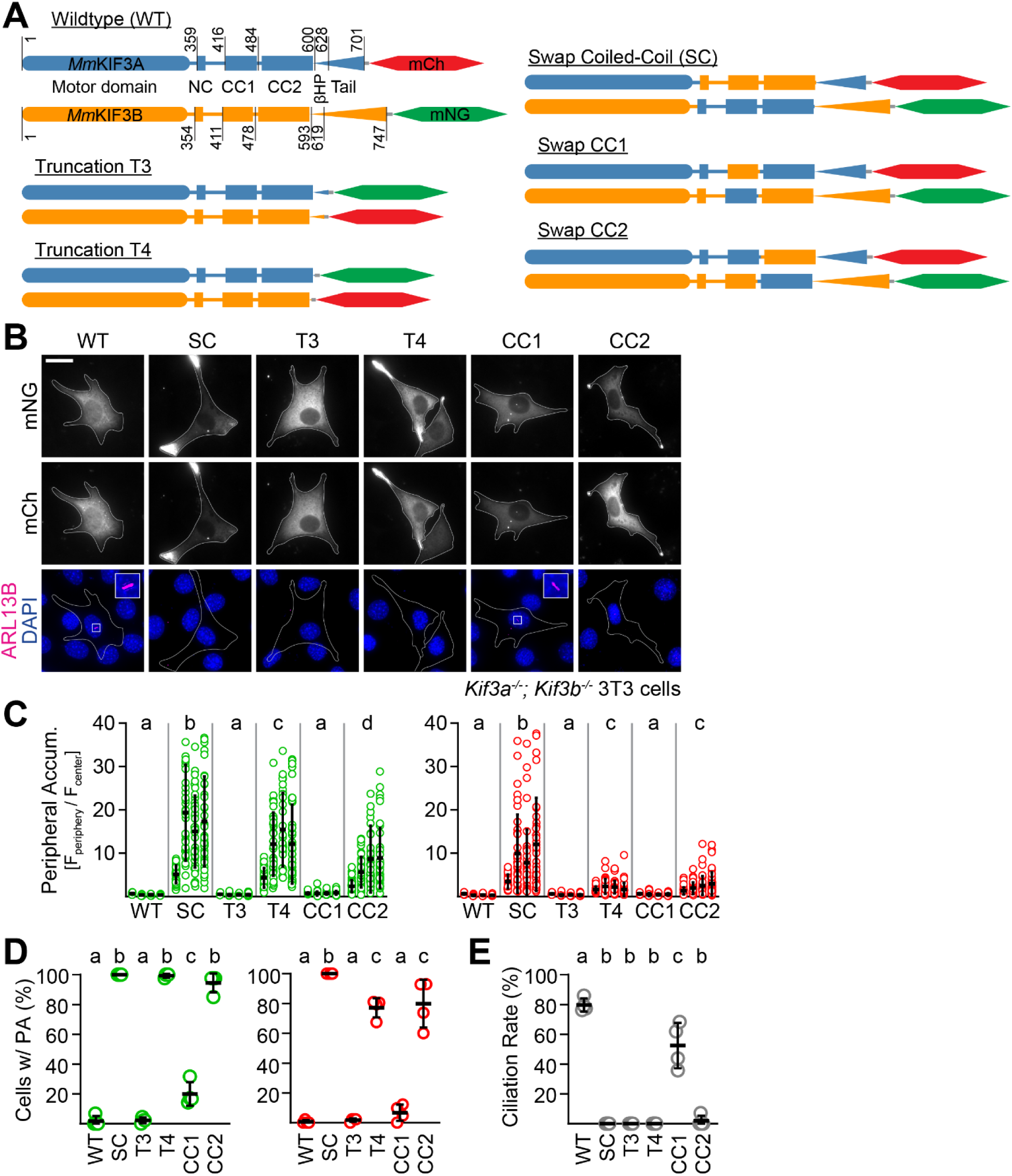
Both the CC2 domain and adjacent proximal tail region are required for autoinhibition. **(A)** Schematic of *Mm*KIF3A (blue) and *Mm*KIF3B (orange) proteins annotated as in Figures 1/2. Amino acid residues at which truncation or splicing occurred to generate the mutant motors are indicated. Full sequences of translated proteins are available in Supplementary Table 1. **(B-E)** *Kif3a^-/-^; Kif3b^-/-^* 3T3 cells were co-transfected with two expression plasmids encoding the pairs of proteins illustrated in panel A, and primary cilium formation was stimulated by serum starvation for 48 hours. **(B)** Representative image of each condition. Cilia were visualized by immunostaining with an antibody against the ciliary membrane marker ARL13B (magenta) and nuclei were visualized with DAPI. Insets show an enlarged region of interest containing the primary cilium. Scale bar = 20 μm. **(C)** Quantification of peripheral accumulation (PA) ratio (fluorescence intensity of a peripheral [F_periphery_] divided by a perinuclear [F_center_] region of interest) in transfected cells in each condition (mNG, green, left; mCh, red, right). **(D)** Quantification of the percentage of cells in each condition with PA ratios greater than 1 (mNG, green, left; mCh, red, right) **(E)** Quantification of the ciliation rate (percentage of transfected cells with a cilium) in each condition. Data represent four biological replicates, with 34-44 cells analyzed per condition per replicate. Data are presented as mean ± SD and were analyzed via one-way ANOVA followed by Tukey’s post hoc tests. Groups that do not share a letter differ significantly (compact letter display). All summary statistics, ANOVA outputs, and Tukey post hoc results are provided in Supplementary Tables 2-4.

As in previous experiments, wildtype (WT) motors expressed in *Kif3a^-/-^; Kif3b^-/-^* cells showed minimal peripheral accumulation, whereas Swap Coil (SC) motors robustly accumulated at the cell periphery (**Fig. 4B-D**). T3 truncation mutants were indistinguishable from WT in both the extent of peripheral accumulation and the fraction of cells exhibiting this phenotype, consistent with results obtained with N-terminally tagged motors (see **Fig. 2**). In contrast, complete removal of the tail domains in the T4 constructs led to strong peripheral accumulation in most expressing cells. Peripheral accumulation appears more prominent in the mNG channel than in the mCh channel, which could be due to unequal expression levels of the constructs or unequal propensity to form homodimers (Funabashi et al., 2018; Jiang et al., 2026), but we do not attribute any biologically meaningful significance to this difference. Strikingly, C-terminal tagging resulted in a markedly stronger peripheral accumulation than the corresponding N-terminally tagged truncation mutants described earlier, highlighting the importance of tag position. We interpret these results with caution, as C-terminal tagging has been reported to promote kinesin activation (Hammond et al., 2010). Nevertheless, the selective motor activation observed with T4, but not T3, truncation supports the conclusion that the proximal tail regions adjacent to CC2, including the β-hairpin motifs, contribute substantially to autoinhibition.

Constructs in which the CC1 domains were swapped exhibited a mild phenotype, with a slight increase in peripheral accumulation that was generally not distinguishable from wildtype (**Fig. 4B-D**). However, a significant fraction of cells exceeded the accumulation threshold as measured by mNG levels, suggesting subtle yet significant activation (**Fig. 4D**). In contrast, Swap CC2 motors were robustly activated, with strong peripheral accumulation and a high proportion of cells exhibiting this phenotype (**Fig. 4B-D**). Interestingly, although both the T4 truncations and Swap CC2 constructs showed strong activation phenotypes, neither accumulated to the same degree as the Swap Coiled-coil (SC) motors, suggesting that the CC2 and proximal tail domains are both required for full autoinhibition. Consistent with the results above, loss of autoinhibition with the Swap constructs inversely correlated with the ability of the motors to support ciliogenesis (**Fig. 4E**). Interpretation of the T3 and T4 tail truncation phenotypes is, however, complicated by the likelihood that these truncations also impair binding to the accessory subunit KAP3 and/or IFT cargo (Funabashi et al., 2018; Webb et al., 2025).

### The motor domain of KIF3A is more important than that of KIF3B for autoinhibition and ciliogenesis

Having implicated the motor domains as well as CC2 and the proximal tail regions in autoinhibition, we next asked whether the two motor subunits contribute equally to this mechanism. To address this question, we combined Swap Motor (SM) and wildtype (WT) constructs to generate heterodimers containing either two KIF3A motor domains (AA Motor) or two KIF3B motor domains (BB Motor) (**Fig. 5A**), analogous to a strategy used previously (Brunnbauer et al., 2010).

**Figure 5.**
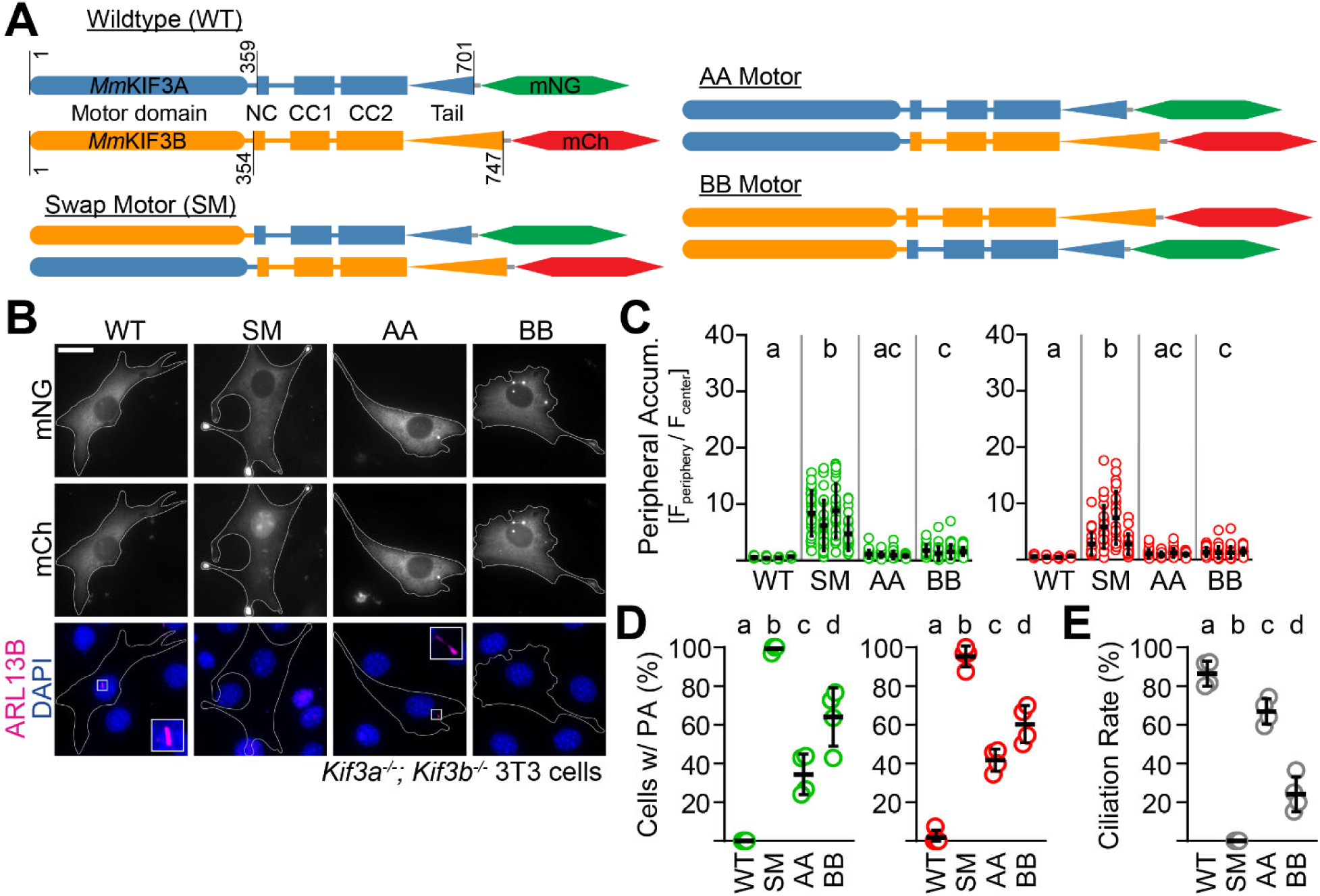
KIF3A and KIF3B motor domains can drive IFT along axonemes and are required for regulating autoinhibition, but the KIF3A motor domain is more important. **(A)** Schematic of *Mm*KIF3A (blue) and *Mm*KIF3B (orange) proteins annotated as in Figure 1. Amino acid residues at which splicing occurred to generate the mutant motors are indicated. Full sequences of translated proteins are available in Supplementary Table 1. (B-E) *Kif3a^-/-^; Kif3b^-/-^* 3T3 cells were co-transfected with two expression plasmids encoding the pairs of proteins illustrated in panel A, and primary cilium formation was stimulated by serum starvation for 48 hours. **(B)** Representative image of each condition. Cilia were visualized by immunostaining with an antibody against the ciliary membrane marker ARL13B (magenta), and nuclei were visualized with DAPI. Insets show an enlarged region of interest containing the primary cilium. Scale bar = 20 μm. **(C)** Quantification of peripheral accumulation (PA) ratio (fluorescence intensity of a peripheral [F_periphery_] divided by a perinuclear [F_center_] region of interest) in transfected cells in each condition (mNG, green, left; mCh, red, right). **(D)** Quantification of the percentage of cells in each condition with PA ratios greater than 1 (mNG, green, left; mCh, red, right) **(E)** Quantification of the ciliation rate (percentage of transfected cells with a cilium) in each condition. Data represent four biological replicates, with 24-35 cells analyzed per condition per replicate. Data are presented as mean ± SD and were analyzed via one-way ANOVA followed by Tukey’s post hoc tests. Groups that do not share a letter differ significantly (compact letter display). All summary statistics, ANOVA outputs, and Tukey post hoc results are provided in Supplementary Tables 2-4.

Consistent with prior experiments, WT motors showed minimal peripheral accumulation, whereas SM constructs robustly localized to the cell periphery in most expressing cells (**Fig. 5B-D**). AA Motor constructs showed only a mild phenotype, with a slight increase in peripheral accumulation. However, a larger fraction of cells expressing AA Motor constructs showed peripheral accumulation compared to WT, suggesting subtle but significant activation (**Fig. 5D**). By contrast, BB Motor constructs were more strongly activated, with greater peripheral accumulation and a higher proportion of cells displaying this phenotype than either WT or AA Motors (**Fig. 5B-D**). Nevertheless, neither the AA Motor nor the BB Motor constructs produced the robust activation observed with the SM controls. These results indicate that both motor domains participate in autoinhibition, but that the KIF3A motor domain makes the larger contribution. As in the preceding analyses, reduced autoinhibition was associated with a reduced rescue of ciliogenesis (**Fig. 5E**).

### Moderate autoinhibition loss yields impaired motors that strongly accumulate at the cilium tip

Across the preceding experiments, one consistent pattern emerged: constructs with partial loss of autoinhibition retained some ability to support ciliogenesis, but not to wildtype (WT) levels (**Figs. 3E, 4E, 5E**). We therefore asked whether these partially functional motors produce a distinct ciliary phenotype. As expected, ciliogenesis rescue decreased as motor activation increased (**Figs. 6A, 3C-E, 5C-E**). The length of cilia varied widely for all constructs (**Fig. 6B**). In cells expressing Swap Tail (ST) and AA Motor constructs, we observed several unusually long cilia (> 6 μm). Cells expressing the BB Motor constructs had a slightly but significantly reduced average cilium length as compared to WT and ST expressing cells. Notably, whereas wildtype (WT) motors exhibited characteristic diffuse cytoplasmic expression, the chimeric motors were highly likely to form perinuclear motor puncta, defined as fluorescence intensity exceeding twice the surrounding background (**Fig. 6C**). We categorized each cell based on ciliation status, presence of a bright motor punctum and its colocalization with the ciliary membrane marker ARL13B and/or the basal body/centrosome marker pericentrin (PCN). Although we observed some cells in which bright motor puncta did not colocalize with the ciliary marker ARL13B (**Fig. 6D-F**, light yellow category), the majority of motor puncta did (blue and purple categories). Surprisingly, a large number of cells that contained ARL13B-positive bright motor puncta did not have a detectable cilium (light purple and light blue categories, depending PCN colocalization). In ciliated chimeric motor-expressing cells, motor puncta were never observed at the base of the cilium. Rather, they usually formed at the ciliary tip (medium purple category), or occasionally could be observed adjacent to the cilium tip (dark purple category). We proposed that the latter could originate from ciliary tip shedding and tested this hypothesis using live-cell imaging, employing ARL13B-mStayGold as a ciliary membrane marker. To our surprise, in 3-hour-long time-lapse videos, we never observed tip shedding (n=7; **Supplementary Video 1**). Instead, we frequently observed cilia with highly fluctuating lengths, with an overall bias toward shrinking. Shrinking of the cilium could, in theory, result in colocalization of motor puncta with both PCN and ARL13B. However, we rarely observed that phenotype in fixed-cell experiments (**Fig. 6D-E**, light blue category). Future experiments will focus on studying and characterizing this phenomenon. Taken together, these findings strengthen the inverse relationship between kinesin-2 activation and functional rescue, and further suggest that partial loss of autoinhibition yields motors that accumulate at the ciliary tip and are potentially associated with axonemal instability.

**Figure 6.**
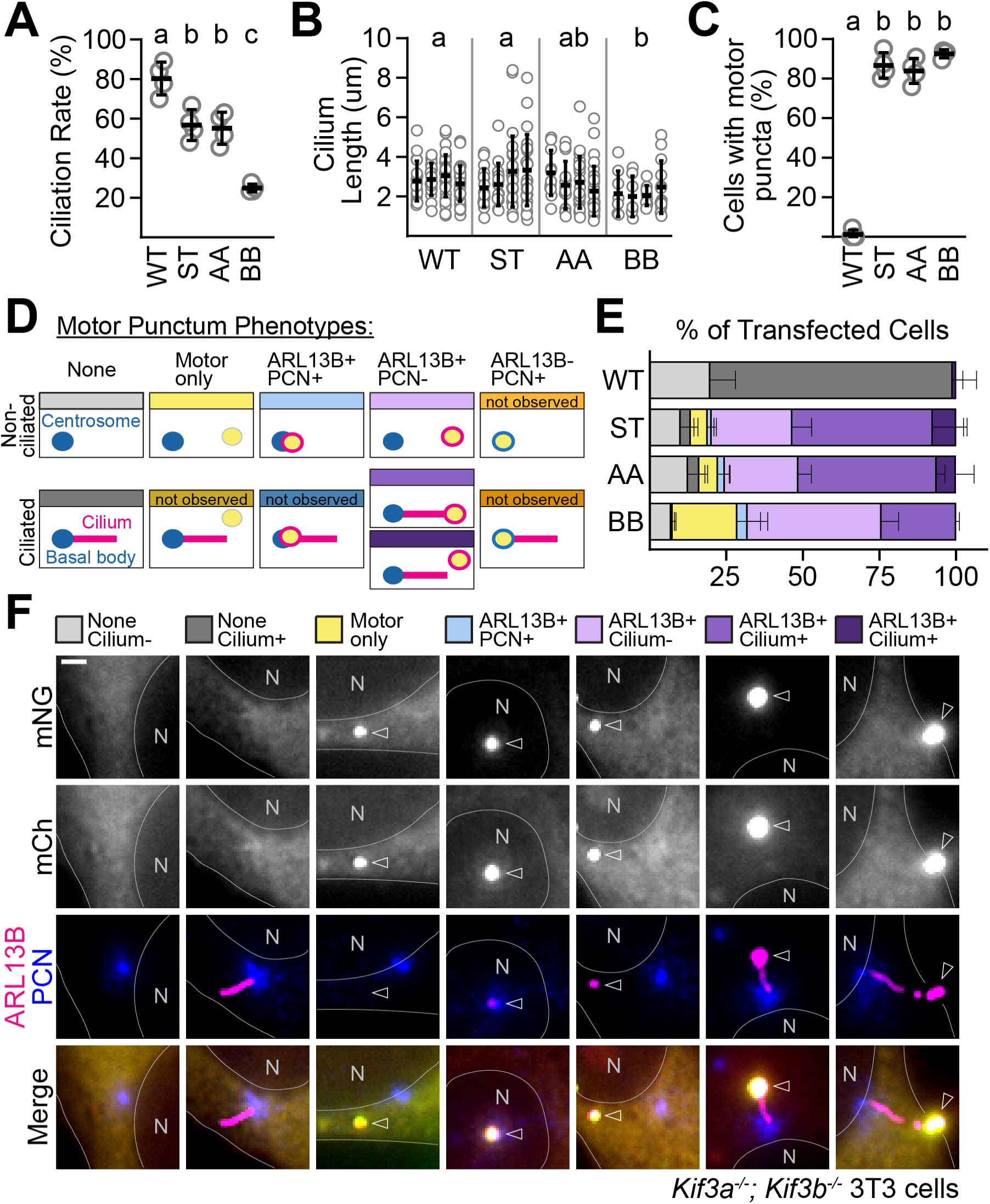
Chimeric KIF3A/KIF3B motors with impaired autoinhibition accumulate at ciliary tips. *Kif3a^-/-^; Kif3b^-/-^* 3T3 cells were co-transfected with two expression plasmids encoding pairs of proteins for the indicated motors, and primary cilium formation was stimulated by serum starvation for 48 hours. The primary cilium and centrosome/basal body were visualized via immunostaining with ARL13B and pericentrin (PCN) antibodies, respectively. **(A)** Quantification of the ciliation rate (percentage of transfected cells with a cilium) in each condition. **(B)** Quantification of cilium lengths as measured by the ARL13B signal. **(C)** Quantification of the percentage of cells in each condition that contained a bright motor punctum. **(D-F)** Each transfected cell analyzed was categorized based on ciliation state, the presence/absence of a bright motor punctum, and its colocalization with ARL13B and/or PCN. **(D)** Definition of each category along with the color key. **(E)** The percentage of analyzed cells in each category in each condition. **(F)** Representative image of one cell from each category. Arrowheads indicate the motor puncta, identified in mNG and mCh. N = nucleus. Scale bar = 2 μm. Data represent four biological replicates, with 27-48 cells analyzed per condition per replicate. Data are presented as mean ± SD and were analyzed via one-way ANOVA followed by Tukey’s post hoc tests. Groups that do not share a letter differ significantly (compact letter display). All summary statistics, ANOVA outputs, and Tukey post hoc results are provided in Supplementary Tables 2-4.

## DISCUSSION

Proteins of the kinesin superfamily are subject to autoinhibition, a property that likely conserves ATP and, more importantly, is essential for physiological motor function. Here, using *Kif3a^−/−^; Kif3b^−/−^* double-knockout mouse embryonic fibroblasts and cell-based assays, we dissect kinesin-2 autoinhibition and its role in ciliogenesis. We chose a cellular system because *in vitro* assays do not fully recapitulate the intracellular environment or allow conclusions about native motor function. Because truncation-based mapping disrupted motor dimerization, we primarily used domain swaps between KIF3A and KIF3B to analyze this heterodimeric motor. This approach confirms that the previously identified β-hairpins in the tail and the CC2 domain act synergistically in autoinhibition (Webb et al., 2025), while our data place greater emphasis on the contribution of the CC2 domain. Notably, our results suggest that the same C-terminal regions that mediate KIF3A/KIF3B heterodimerization also participate in autoinhibitory interactions with the motor domains, linking motor assembly and autoregulation. We also find that both KIF3A and KIF3B motor domains can support movement along the axoneme and mediate ciliogenesis, albeit less efficiently, indicating that a heterodimeric motor-domain complex is not strictly required for IFT. Instead, heterodimerization appears to be required for fine-tuning motor regulation. Consistent with this view, loss of autoinhibition directly correlates with reduced ability to support cilium assembly. Within this framework, we propose that autoinhibition promotes kinesin-2 enrichment at the ciliary base by preventing premature engagement with cytosolic microtubules, which would lead to motor depletion to the cell periphery, and by facilitating motor return from the ciliary tip to the base after transport.

As with kinesin-1, kinesin-2 can adopt either a compact, autoinhibited conformation or an extended, active conformation (Friedman and Vale, 1999; Hammond et al., 2010; Imanishi et al., 2006; Wedaman et al., 1996). Entry into the autoinhibited state has been proposed to depend on a conserved flexible stalk segment containing a diglycine motif. In homodimeric kinesin-2, deletion of this segment or mutation of the diglycine motif activates the motor *in vitro* and *in vivo* (Hammond et al., 2010; Imanishi et al., 2006; Xie et al., 2024), and mutation of the corresponding motif similarly activates heterotrimeric kinesin-2 *in vitro* (Brunnbauer et al., 2010). Consistent with this model, deletion of the entire flexible segment in mammalian KIF3A and KIF3B robustly activated the motor in cells, confirming that conformational flexibility in this region is essential for kinesin-2 autoinhibition. However, rather than serving as a simple hinge for backfolding, recent structural work suggests that heterotrimeric kinesin-2 undergoes a broader structural collapse involving unzipping and separation of large portions of the coiled-coil stalk (Webb et al., 2025; **Fig. 7A-C**).

**Figure 7.**
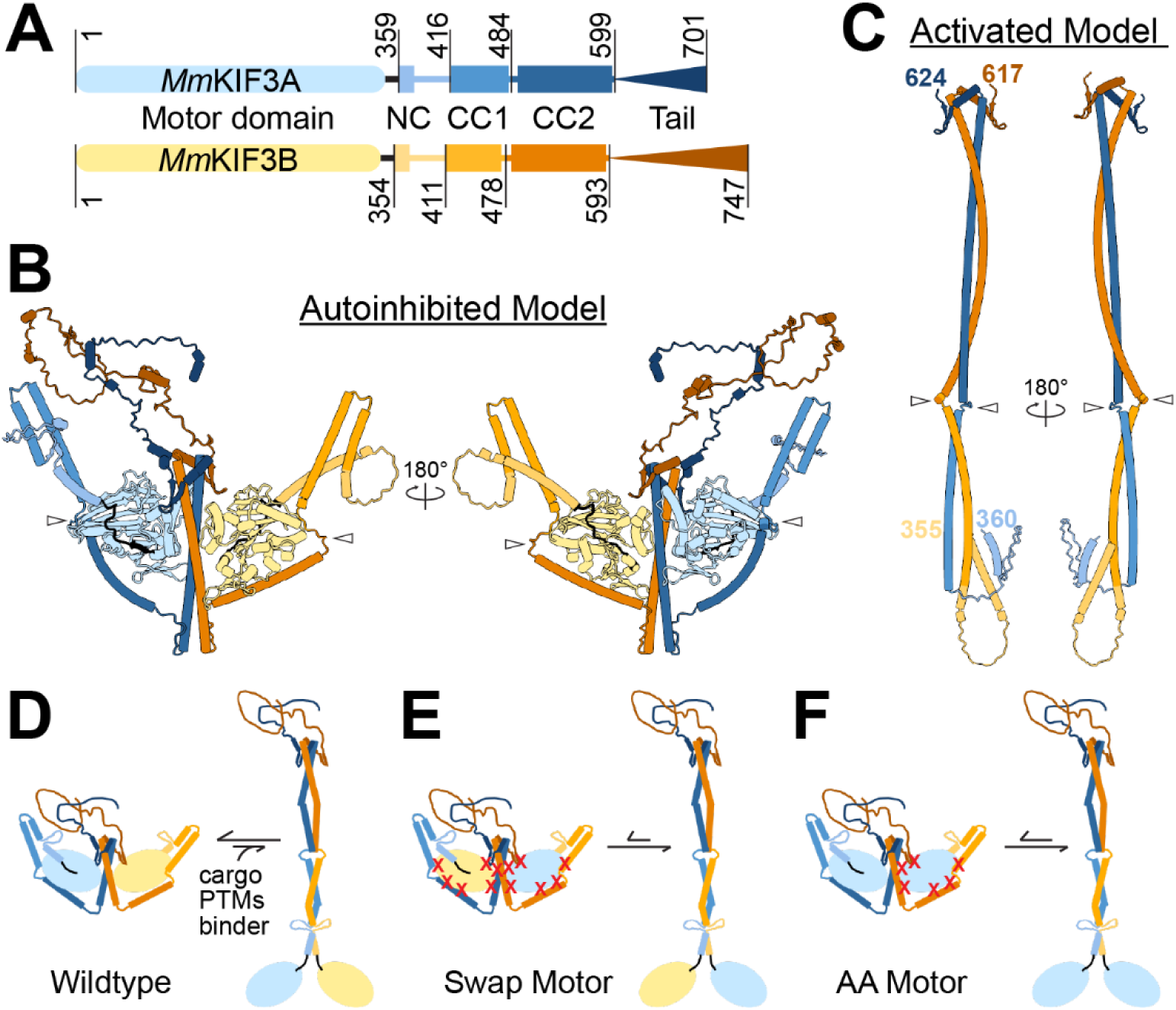
Model of KIF3A/KIF3B autoinhibition and its disruption in the chimeric mutants**. (A)** Schematic of *Mm*KIF3A (blue) and *Mm*KIF3B (orange) proteins annotated as in Figure 1. Amino acid residues at which domain transitions occur are indicated. **(B-C)** AlphaFold3-generated structural predictions of KIF3A/KIF3B. Domains were color-coded according to the schematic from **(A)** using ChimeraX. Open arrowheads indicate the flexible region between CC1 and CC2. **(B)** The predicted autoinhibited structure shows potential sites of interaction between the motor domains and the coiled-coil stalk and adjacent proximal tail region. The structure was generated by including the accessory protein KAP3A (NP_001292572.1) in the AlphaFold3 input, but it has been hidden in the model for clarity. **(C)** The predicted activated structure of the KIF3A/KIF3B stalk, including the neck coils and β-hairpins (*Mm*KIF3A: 360-624; *Mm*KIF3B: 355-617). **(D-F)** Proposed conformational equilibria between autoinhibited and activated states of WT and mutant KIF3A/KIF3B motors. Red X’s indicate likely areas of interaction in wildtype that are disrupted by the domain swaps. **(D)** Wildtype motors are likely to exist mostly in the autoinhibited conformation, but can be activated by cargo, other binders, and/or post-translational modifications (PTMs). **(E)** KIF3A/KIF3B Swap Motors have lost interactions that stabilize the autoinhibited conformation and are more likely to exist in the activated conformation. **(F)** AA motors have lost fewer interactions than the Swap Motors that stabilize the autoinhibited conformation. Thus, the dynamic pool of autoinhibited motors is larger, increasing the availability of motors at the ciliary base.

The specific interactions that stabilize this compact state remain incompletely defined. The motor domains of mouse KIF3A and KIF3B are highly conserved, making it striking that swapping them is sufficient to disrupt autoinhibition. Prior efforts to define the independent contributions of the two motor domains to motility and autoinhibition have yielded mixed conclusions. *In vitro* studies of “AA Motor” and “BB Motor” chimeras found both motors to be processive, with only modest differences in velocity (Albracht et al., 2016; Guzik-Lendrum et al., 2015; Zhang and Hancock, 2004). By contrast, work on the *C. elegans* heterotrimeric kinesin-2 motor KLP11/KLP20/KAP1 suggested that the KLP11 motor domain, the KIF3B homolog, is nonprocessive but contributes asymmetric autoinhibition of its processive partner KLP20, the KIF3A homolog (Brunnbauer et al., 2010). Our cell-based data indicate that both KIF3A and KIF3B motor domains can support motility and IFT along axonemal microtubules. At the same time, they suggest that the motor-domain contributions to autoinhibition are also asymmetric, with KIF3A making the stronger contribution, since the BB Motor is more active than the AA Motor. Because kinesin-2 serves distinct roles across species, and because the *C. elegans* heterotrimeric motor acts only as an accessory IFT motor, some of these differences may reflect task-specific tuning.

Recent work identified a conserved tail β-hairpin motif in the KIF3A and KIF3B tail domains and, using *in vitro* assays, showed that this motif is a key determinant of autoinhibition (Webb et al., 2025). Our data support this model and extend it by showing that the β-hairpin also contributes to kinesin-2 regulation in cells. However, our results add an important nuance: robust activation of truncated motors was observed only when the constructs carried a C-terminal fluorescent protein tag, potentially sensitizing the motor for activation. Additionally, the failure of partial tail truncations to strongly activate the motor suggests that sequences distal to the β-hairpin are likely dispensable for autoinhibition. Webb *et al*. proposed that negatively charged residues at the apex of the β-hairpins interact electrostatically with the motor domains. Because the overall β-hairpin structure, the negatively charged apex, and the corresponding positively charged residues in the motor domain are conserved between KIF3A and KIF3B, these interactions are likely preserved in the Swap Tail mutant, potentially explaining its relatively mild phenotype. However, residues surrounding the charged apex are less well conserved, and the Swap Tail mutant is still activated. Together, these observations suggest that the charge-based contacts identified by Webb et al. are insufficient to fully account for the tail domain’s contribution to autoinhibition.

Our data similarly refine the current model of CC2-dependent regulation. Webb *et al*. proposed, based on structural data and predictive modeling, that the motor domains interact electrostatically with the C-terminal coiled-coil region, and found that mutation of six negatively charged residues in CC2 activated the motor *in vitro* (Webb et al., 2025). However, these residues are identical in the KIF3A and KIF3B heptad repeat structure. If those contacts were sufficient to explain autoinhibition, the Swap Coil mutant would not be expected to show a strong phenotype. Instead, Swap Coil phenocopies Swap Motor, with both showing similarly strong peripheral accumulation that exceeds that of any other mutant tested. Together, these findings suggest that motor-coiled-coil electrostatic interactions are important, but the autoinhibitory interface extends beyond the six CC2 residues previously identified. Rather, our results favor a broader interaction surface involving additional contacts, as schematically illustrated in **Fig. 7D-F**.

Our experiments also inform our understanding of how kinesin-2 assembles. KIF3A/KIF3B/KAP3 is generally thought to form a heterotrimer in which KIF3A and KIF3B heterodimerize through their stalks, and KAP3 binds the tails (Jiang et al., 2025; Wedaman et al., 1996; Yamazaki et al., 1995). Multiple truncation-based studies concluded that the C-terminal portion of the stalk acts as a dimerization seed (De Marco et al., 2001; Funabashi et al., 2018; Vukajlovic et al., 2011), and our results are consistent with this view. However, they also suggest that determinants of stable heterodimer formation are not confined to CC2. Instead, our data support a broader C-terminal interaction interface that includes tail sequences adjacent to CC2, including the recently described β-hairpins. This interpretation is reinforced by recent cryo-EM structures and AlphaFold-based models of the C-terminal regions of the mouse and *C. elegans* heterotrimeric kinesin-2, which predict extensive contacts within CC2 and interactions between CC2 and neighboring tail sequences (Jiang et al., 2025; Ren et al., 2025). Together with our autoinhibition data, these findings suggest that the same structural elements that stabilize KIF3A/KIF3B heterodimerization also engage the motor domains to stabilize the autoinhibited state, thereby coupling motor assembly to autoregulatory control.

Together, our findings support a model in which kinesin-2 autoinhibition is not only an energy-conserving mechanism but also a central determinant of motor function during IFT (Imanishi et al., 2006; Webb et al., 2025). Using swap mutants that are unlikely to strongly disrupt IFT-train binding or processive motility, we show that graded loss of autoinhibition correlates directly with impaired ciliogenesis and likely reflects a shift in the equilibrium between autoinhibited and active conformations, as illustrated in **Fig. 7D-F**. We propose that the swap mutations primarily impair ciliogenesis by destabilizing the autoinhibited state, thereby promoting premature microtubule engagement. In nondividing fibroblasts, where the ciliary base functions as a microtubule-organizing center, such premature engagement would be expected to drive motors away from the ciliary base, reducing their availability for anterograde IFT initiation. This model is consistent with the exclusion of constitutively active OSM-3 from *C. elegans* cilia (Xie et al., 2024). Furthermore, this model explains the partial functionality of moderately activated motors, relative to the loss of function observed in more strongly activated mutants. The accumulation of moderately activated motors at the ciliary tip further supports the model in **Fig. 7**, as these motors would be less likely to re-enter the autoinhibited state after transport and therefore less likely to recycle efficiently to the base. More broadly, these findings align with growing evidence that inappropriate kinesin activation is deleterious to cells and causes disease (Baron et al., 2022; Bianchi et al., 2016; Cheng et al., 2014; Chiba et al., 2019; Cong et al., 2021), and suggest that tight autoregulatory control of kinesin-2 is essential for both cilium assembly and ciliary homeostasis.

## METHODS

### Plasmids

Expression plasmids were generated using the pEGFP-N1 vector backbone (Clontech) via established molecular cloning techniques. These include conventional PCR, splice by overlap extension (Horton et al., 1989), and commercial DNA synthesis (Twist Bioscience), followed by restriction digestion. Mouse KIF3A (NP_032469.2), mouse KIF3B (NP_032470.3), mCherry, and mNeonGreen (Allele Biotechnologies) were codon-optimized for mammalian expression. The translated protein sequence for each expression plasmid used is listed in Supplementary Table 1. Plasmid sequences were verified by Sanger sequencing (Eurofins) or by long-read whole-plasmid sequencing (Plasmidsaurus).

### Cell Culture

*Kif3a^-/-^; Kif3b^-/-^* mouse embryonic fibroblast 3T3 cells (Engelke et al., 2019) and COS-7 cells (ATCC Cat# CRL-1651) were cultured in D-MEM (Corning Cat# 15017CV) supplemented with 10% Hyclone FetalClone III serum (Cytiva Cat# SH30109.03) and 4 mM L-glutamine (Alfa Aesar Cat# J6057322) at 37°C and 5% CO_2_.

For peripheral accumulation and ciliogenesis rescue experiments, cells were seeded at a density of 1×10^5^ cells/well on glass coverslips (Electron Microscopy Sciences Cat# 72229-01) inserted into wells of a 12-well plate (Fisher Scientific Cat# FB012928). Approximately 16 hours later, the media was switched to starvation media (1% FetalClone III), and 800 ng total plasmid DNA per well was transfected. All transfections were performed with Lipofectamine 2000 (Thermo Fisher Scientific Cat# 11668019) according to the manufacturer’s protocol. 48 hours following transfection, cells were fixed and processed for immunofluorescence (see below).

For the VIP assay, COS-7 cells were seeded in a 6-well plate (Corning Cat# 3516) at a density of 3.6×10^5^ cells/well. 5–6 hours later, fresh media were applied, and 1.6 μg total plasmid DNA per well was transfected as above. Approximately 16 hours after transfection, lysates were collected and analyzed (see below).

### Immunofluorescence

Cells were fixed with 10% buffered formalin (Fisher Scientific Cat# 11-002-205) for 15 min, and immunostaining was performed as previously described (Engelke et al., 2019). Primary antibodies used were: rabbit polyclonal anti-ARL13B (Proteintech Cat# 17711-1-AP, RRID:AB_2060867; 1:1000), mouse monoclonal anti-ARL13B (N295B/66) (Neuromab Cat# 75-287, RRID:AB_2877361; 1:200); rabbit polyclonal anti-pericentrin (Abcam Cat# ab4448, RRID: AB_304461; 1:1000). Secondary antibodies used were: goat anti-rabbit-AlexaFluor 647 (Thermo Fisher Scientific Cat# A-21245, RRID:AB 2535813; 1:500); goat anti-mouse-AlexaFluor 647 (Thermo Fisher Scientific Cat# A-21236, RRID:AB 2535805; 1:500) goat anti-rabbit-AlexaFluor 405 Plus (Thermo Fisher Scientific Cat# A-48255, RRID:AB 2890536; 1:500). Nuclei were visualized with DAPI (Biotium Cat# 40043; 1:10,000). Coverslips were mounted onto microscope slides using Prolong Gold (Thermo Fisher Scientific Cat# P36930) (Fisher Scientific Cat# 12–544-7).

### Microscopy and analysis

Images for Fig. 1 and 3-6 were acquired using an inverted, semi-automated BZ-X810 epifluorescence microscope (Keyence) equipped with a Plan Apochromat 40×/0.95 objective (16-bit acquisition). Images for Fig. 2 were acquired using a Nikon Ti2-E inverted motorized microscope equipped with a CFI Plan Apo λD 40×/0.95 objective and an ORCA-Fusion BT sCMOS camera (12-bit acquisition). Exposure times and illumination power settings, when applicable, were optimized for each channel and then kept constant across conditions within each experiment, while all other camera settings were identical across the study. Images were analyzed with ImageJ software enhanced by the FIJI package (Schindelin et al., 2012). Ciliation rates were calculated as the percentage of transfected cells with a cilium ≥0.5 μm. Cilium lengths were measured manually. Peripheral accumulation was assayed by measuring mean fluorescence intensity in a small circular ROI (Fig. 1, 3-5: 0.43 μm^2^; Fig. 2: 0.49 μm^2^) at the cell periphery (F_periphery_) and another next to the nucleus (F_center_). The accumulation ratio was calculated as F_periphery_/F_center_. For characterization of motor puncta, an ROI was manually drawn around each punctum (fluorescence intensity>2X cytoplasmic background in mNG and mCh), evaluated for signal in the PCN and ARL13B channels, and categorized manually. Cilia and ROI from all images have been non-destructively annotated in the dataset published with this study. All experiments were performed in four biological replicates, and 20-50 cells per condition per replicate were analyzed.

### VIP Assay

Crude whole-cell lysates were collected according to a previously published protocol (Engelke et al., 2016). The modified VIP assay (Katoh et al., 2018) was performed as previously described (Adams et al., 2024). Beads were imaged with a Keyence epifluorescence microscope and a PlanFluor 10×/0.30 objective, with exposure times optimized for each channel and identical camera settings across conditions. For each bead in one field of view, an ROI encompassing the entire bead was drawn, fluorescence intensity was measured for mCherry and mNG, and the ratio F_mCh_/F_mNG_ was calculated. For each bead, this ratio was normalized to the wildtype average for that experiment, and then the average normalized value across all beads in each condition was computed. All images have been non-destructively annotated in the dataset published with this study. All experiments were performed in three biological replicates, and 30-60 beads per condition per replicate were analyzed.

### Statistics

Prism 11.0.0 (GraphPad) was used to graph the data and perform statistical analyses. Data are displayed as mean ± SD and were analyzed by one-way ANOVA followed by Tukey’s post hoc tests. Statistical significance is indicated using a compact letter display. All experiments were performed in 3-4 independent biological replicates. The number of cells analyzed per group (n) and summary statistics (mean ± SD) for each experiment are provided in Supplementary Table 2. Full ANOVA results, including F statistics and p-values, are provided in Supplementary Table 3. The results of the Tukey post hoc tests are provided in Supplementary Table 4.

### Other software

AlphaFold3-generated structural predictions were displayed using ChimeraX 1.9 (Meng et al., 2023).

### Use of artificial intelligence

AlphaFold3 (Abramson et al., 2024) was used to predict the structure of heterotrimeric kinesin-2, and models of those structures, with KAP3 protein chains hidden, were displayed.

OpenAI’s GPT-5.4 was used to refine the draft text, making it more concise and improving its scientific style.

## ACKNOWLEDGEMENTS

This study was supported by the National Institute of General Medical Sciences of the National Institutes of Health under award number R35GM147641 to MFE. The content is solely the authors’ responsibility and does not necessarily represent the official views of the National Institutes of Health. We would also like to thank Dr. Tongye Shen (University of Tennessee, Knoxville) for helpful discussions.

## SUPPLEMENTARY MATERIAL

### Supplementary Methods

**Time-lapse imaging.** *Kif3a^-/-^; Kif3b^-/-^* 3T3 cells were seeded on glass-bottom 35 mm dishes at a density of 2.4 x 10^5^ cells per dish. Approximately 6 hours later, cells were co-transfected with two expression vectors encoding the AA Motor fused to mCherry as well as *Hs*ARL13B-mStayGold to visualize the ciliary membrane. Following serum starvation for ∼40 hours, the medium was replaced with FluoroBrite DMEM live-imaging media (Gibco Cat# A1896701) supplemented with 1% FetalClone III and 4 mM L-glutamine. SPY650-tubulin (Cytoskeleton Inc Cat# CY-SC503) was added to the medium to visualize the ciliary axoneme. Dishes were maintained at 37°C and 5% CO_2_ using a Tokai Hit warming box during time-lapse imaging. Images were acquired every 1 minute for 182 minutes using the Perfect Focus System on a Nikon Ti2-E inverted motorized microscope equipped with a CFI Plan Apo λD 60×/1.42 oil-immersion objective and an ORCA-Fusion BT sCMOS camera (12-bit acquisition). Image sequences were processed and assembled using FIJI (ImageJ). Frames 101-102, 135-138, and 173-179 were removed due to the cilium going out of focus. Nonlinear adjustments to brightness and contrast were applied to each channel using the FIJI “Enhance Contrast” tool. Text annotations and final video rendering were performed using Adobe Photoshop.

**Supplementary Video 1.** Time-lapse images of a representative cilium in *Kif3a^-/-^; Kif3b^-/-^*3T3 cells expressing the AA Motor (visualized with mCherry, right panel) and ARL13B-mStayGold (mSG) to visualize the ciliary membrane (left panel). SPY650-tubulin was added to the imaging media to visualize the ciliary axoneme (center panel). The initial cilium displays a strong accumulation of motor and ARL13B at the tip. Over the course of 3 hours, the cilium is seen to shrink, and the bulbous ARL13B tip accumulation appears to be resorbed into the ciliary shaft. The motor accumulation fades, likely due to photobleaching. The arrow points to the ciliary base. Scale bar = 5 μm.

## Notes

### Competing Interest Statement

The authors have declared no competing interest.

